# The influence of rare variants in circulating metabolic biomarkers

**DOI:** 10.1101/480699

**Authors:** Fernando Riveros-Mckay, Clare Oliver-Williams, Savita Karthikeyan, Klaudia Walter, Kousik Kundu, Willem H Ouwehand, David Roberts, Emanuele Di Angelantonio, Nicole Soranzo, John Danesh, Eleanor Wheeler, Eleftheria Zeggini, Adam S Butterworth, Inês Barroso

## Abstract

Circulating metabolite levels are biomarkers for cardiovascular disease (CVD). We tested association between rare sequence variants and 226 serum lipoproteins, lipids and amino acids in 7,142 healthy participants. Gene-based association analyses identified novel gene-trait associations with *ACSL1*, *MYCN*, *FBXO36* and *B4GALNT3* (*p*<2.5 × 10^−6^), and confirmed established associations. Regulation of the pyruvate dehydrogenase (PDH) complex was associated for the first time, in gene set analyses, with IDL and LDL parameters, as well as circulating cholesterol (*p*_*METASKAT*_ <2.41 × 10^−6^). Individuals at the lower tails of the distributions of four out of 49 lipoproteins and lipids had an excess of predicted deleterious variants in lipoprotein disorder and metabolism gene sets (*p*_*permutation*_<0.00037). These four traits were CVD risk factors (e.g. S-VLDL-C), demonstrating that rare “protective” variation is a significant contributor to lipoprotein levels in a healthy population. In conclusion, rare variant analysis of these important metabolic biomarkers reveals novel loci and pathways involved in their regulation.

## Introduction

Metabolic measurements reflect an individual’s endogenous biochemical processes and environmental exposures [1,2]. Many circulating lipids, lipoproteins and metabolites have been previously implicated in the development of cardiovascular disease (CVD) [3-6] or used as biomarkers for disease diagnosis or prognosis [7,8]. Understanding the genetic influence on circulating levels of these metabolic biomarkers can help us gain insight into the biological processes regulating these traits, lead to improved aetiological understanding of CVD and identify novel potential therapeutic drug targets. Notable examples of candidate drug targets with support from human genetics are *LDLR* [9,10], *APOB* [11,12] and *PCSK9* [13,14].

Genome-wide association studies (GWAS) focusing on traditionally measured lipid traits have greatly expanded our knowledge into lipid biology and to date more than 250 loci have been robustly associated with total cholesterol (TC), high-density lipoprotein cholesterol (HDL-C), low-density lipoprotein cholesterol (LDL-C), and/or triglycerides (TG) [15-23]. In addition to this, more detailed metabolic profiling using high resolution nuclear magnetic resonance (NMR) measurements has proven helpful to find additional lipid and small molecule metabolism-associated loci with smaller sample sizes, and to assess pleiotropic effects of previously established loci [24-26]. An example of this, is a novel link between the *LPA* locus and very-low-density lipoprotein (VLDL) metabolism (measured by high resolution NMR), with effect sizes twice as large as those found for traditionally measured lipid traits like LDL-C and TC, suggesting these measurements are better at capturing underlying biological processes in lipid metabolism than traditionally measured lipid traits [25]. In this same study, by constructing a genetic risk score using variants associated with lipoprotein(a) levels and using a Mendelian randomisation approach the authors were able to determine a causal link between increased lipoprotein(a) levels on overall lipoprotein metabolism [25].

Despite the at scale usage of exome arrays to capture low-frequency and rare coding variation contributing to lipid and amino acid metabolism [19-22,26], large-scale sequencing studies have the added value of assessing rare variation at single nucleotide resolution across the whole genome, or exome, including the detection of private variants which could have large effects on protein function. These approaches enabled, for example, the discovery of inactivating variants in key proteins which are models for drug target antagonism [27,28].

In this study, we examined the contribution of rare variation (MAF <1%) to 226 serum metabolic measurements in 3,741 individuals with whole-exome sequence (WES) data and 3,420 individuals with whole-genome sequence (WGS) data from the INTERVAL cohort, which consists of healthy blood donors residing in the UK. We identified new gene-lipid trait associations at *ACSL1*, *MYCN*, *FBXO36* and *B4GALNT3* (gene-based analysis *p*< 2.5 × 10^−6^), and linked loss-of-function variation in genes involved in regulation of the pyruvate dehydrogenase (PDH) complex to intermediate-density lipoprotein (IDL) and low-density lipoprotein (LDL) metabolism and circulating cholesterol for the first time (gene-set-based analysis *p* < 2.4 × 10^−6^). We further show that genes near established HDL-C GWAS index single nucleotide polymorphisms (SNPs) are enriched in rare variants associated with a subset of relevant lipid or lipoprotein traits, suggesting this gene set is enriched in likely effector transcripts (i.e. transcripts/genes likely to be causal of the original association) which are also modulated by rare coding variants. Finally, we show enrichment of rare computationally predicted deleterious variation affecting genes involved in lipoprotein metabolism in the lower tails of the phenotype distribution for four lipoprotein subpopulations (cholesterol and esterified cholesterol in small very-low-density-lipoprotein (VLDL), concentration of very small VLDL particles and concentration of small HDL particles) providing evidence for the role of rare coding variants in conferring very low levels of these traits. Overall, the study demonstrates the value of rare variant association studies to uncover novel loci, pathways and possible mechanisms influencing these important disease biomarkers.

## Results

### Gene-based analyses

The sample size of this study gave us limited power to detect novel single variant associations at rare loci (power 9.7% to find an association at *p*<5 × 10^−8^ with MAF 0.1% and beta=1.1), but was well-powered for common variant analysis (power 86.41% to find an association at *p*<5 × 10^−8^ with MAF 1% and beta=0.55). Therefore, single point analyses only confirmed established associations at 34 unique loci (data not shown).

We then sought to discover new gene-trait associations for 226 NMR metabolic biomarkers using rare-variant aggregate tests. For this analysis we used WES data from 3,741 healthy blood donors from the INTERVAL cohort as a discovery dataset (**Methods**). We performed two nested approaches to group rare variants; first just loss-of-function (LoF) variants and secondly, LOF variants plus variants predicted to be likely deleterious by M-CAP (M-CAP score >0.025) [29] (MCAP+LoF) (**Methods**). Genes were taken forward for validation if they reached an arbitrary threshold of *p* <5 × 10^−3^ in the discovery dataset (**Supplementary Tables 1-2**). As previously suggested, to boost power we adjusted for correlated metabolic biomarkers [30,31]. However, to minimise the possible collider bias this could incur, we only did this at the validation stage. This was to ensure there was at least suggestive evidence for association in the discovery stage without adjusting for any metabolite (**Methods**). Validation was performed using whole-genome sequence (WGS) data from 3,401 independent individuals from the same cohort, and we report results from meta-analysis of discovery plus validation datasets that meet conventional gene-level significance (*p*< 2.5 × 10^−6^, **Table 1, Methods**). After meta-analysis, five genes (*APOB*, *APOC3*, *PCSK9*, *PAH*, *HAL*) were associated with 92 different traits with *p*<1.32 × 10^−7^, which is the stringent significance threshold after correcting for the effective number of tested phenotypes (**Table 1, Methods**). All five have been previously associated with their respective traits [24,32,33]. As expected, we found that there is a significant increase in the strength of the association signal for traits when we used other correlated traits as covariates compared to the unadjusted tests [30,31], with the most notable example being a >30 order of magnitude increase in association strength for *PAH* and phenylalanine (**Supplementary Tables 1-2, Table 1**). In total, 32 of the known gene-trait associations met our stringent significance threshold (*p*<1.32 × 10^−7^) only after adjusting for correlated traits (**Supplementary Tables 1-2**).

**Table 1-.** Genes significantly associated (*p*<2.5 × 10^−6^) with at least one trait in gene-based analyses focusing on loss-of-function (LoF) or predicted deleterious missense by M-CAP plus loss-of-function (MCAP+LoF). Genes that meet gene-level significance after adjusting for multiple phenotypes (*p*<1.32 × 10^−7^) are highlighted in bold. Top trait: trait with the smallest p-value after meta-analysis adjusting for correlated metabolites. p-value (covs): p-value of meta-analysis (WES+WGS) after adjusting for correlated metabolites for top trait. If NA, this analysis was not performed for this trait due to no metabolic biomarkers meeting the criteria to be included as covariates in meta-analysis. p-value (raw): p-value of meta-analysis without adjusting for correlated metabolites for top trait. N WES: number of tested variants in WES. N WGS: number of tested variants in WGS. N overlap: number of variants present in both WES and WGS. N traits associated: number of traits that meet gene-wide significance after adjusting for multiple phenotypes (*p*<1.32 × 10^−7^), genes meeting standard gene-wide significance (2.5 × 10^−6^) in parenthesis. Driven by single variant?: Yes if after conditioning on top associated variant the meta-analysis association disappears (*p*>0.05). IDL-TG: Triglycerides in IDL. XS-VLDL-TG: Triglycerides in very small VLDL. Phe: Phenylalanine. His: Histidine. IDL-FC: Free cholesterol in IDL. IDL-P: Concentration of IDL particles. M-VLDL-L: Total lipids in medium VLDL. Gly:Glycine. XL-HDL-FC: Free cholesterol in very large HDL. IDL-CE %: Cholesterol esters to total lipids ratio in IDL. L-VLDL-FC %: Free cholesterol to total lipids ratio in large VLDL. XXL-VLDL-C %: Total cholesterol to total lipids ratio in extremely large VLDL.

| LoF |  |  |  |  |  |  |  |  |
| --- | --- | --- | --- | --- | --- | --- | --- | --- |
| Gene | Top trait | p-value (covs) | p-value (raw) | N WES | N WGS | N overlap | N traits associated | Driven by single variant? |
| *APOB* | IDL-TG | $3.20 \times 10^{-13}$ | $1.72 \times 10^{-10}$ | 6 | 5 | 0 | 45 (57) | No |
| *APOC3* | XS-VLDL-TG | $6.10 \times 10^{-13}$ | $3.58 \times 10^{-12}$ | 3 | 2 | 2 | 46 (56) | No |
| *PAH* | Phe | $5.82 \times 10^{-11}$ | $8.25 \times 10^{-3}$ | 4 | 3 | 1 | 1 (1) | Yes |
| MCAP+LoF |  |  |  |  |  |  |  |  |
| Gene | Top trait | p-value (covs) | p-value (raw) | N WES | N WGS | N overlap | N traits associated | Driven by single variant? |
| *PAH* | Phe | $8.33 \times 10^{-63}$ | $1.67 \times 10^{-28}$ | 13 | 17 | 6 | 1 (1) | No |
| *HAL* | His | NA | $3.72 \times 10^{-42}$ | 9 | 10 | 5 | 1 (1) | No |
| *APOC3* | XS-VLDL-TG | $5.46 \times 10^{-11}$ | $2.15 \times 10^{-10}$ | 1 | 2 | 1 | 26 (40) | No |
| *PCSK9* | IDL-FC | $2.39 \times 10^{-10}$ | $1.11 \times 10^{-7}$ | 3 | 3 | 1 | 29 (34) | No |
| *ACSL1* | IDL-P | $1.82 \times 10^{-7}$ | $1.76 \times 10^{-4}$ | 4 | 6 | 2 | 0 (1) | Yes |
| *MYCN* | M-VLDL-L | $6.20 \times 10^{-7}$ | $3.97 \times 10^{-6}$ | 7 | 8 | 3 | 0 (5) | No |
| *ALDH1L1* | Gly | NA | $4.56 \times 10^{-7}$ | 33 | 38 | 19 | 0 (1) | No |
| *SCARB1* | XL-HDL-FC | NA | $6.93 \times 10^{-7}$ | 24 | 18 | 10 | 0 (6) | No |
| *FBXO36* | IDL-CE % | NA | $1.98 \times 10^{-6}$ | 5 | 2 | 1 | 0 (1) | Yes |
| *B4GALNT3* | L-VLDL-FC % | NA | $7.59 \times 10^{-7}$ | 27 | 22 | 13 | 0 (1) | No |
| *LIPC* | XXL-VLDL-C % | NA | $9.03 \times 10^{-7}$ | 27 | 29 | 11 | 0 (2) | No |

In addition, we found 15 gene-trait associations in seven genes meeting standard gene-level significance before adjusting for multiple traits (*p*< 2.5 × 10^−6^) which also had nominal evidence of association in the validation cohort (*p*< 0.05). Nine of these were gene-trait associations in three established genes (*ALDH1L1*, *SCARB1*, *LIPC*, **Table 1**), suggesting that other results achieving this significance threshold may warrant being prioritised for additional follow-up to establish their validity. In particular amongst the remaining four genes, the association between IDL particle concentration (IDL-P) and *ACSL1* (*p* = 1.82 × 10^−7^), as well as, the associations of multiple very-low-density lipoprotein (VLDL) traits to *MYCN* (min *p* = 6.20 × 10^−7^) merit further exploration as both genes have been previously linked to lipid metabolism in mouse studies [34-36].

### Gene set analyses

To find links between predicted loss-of-function rare variants and metabolic biomarker biology, we next explored associations of these variants in 7,150 gene sets. To this end, we used two biological pathway databases (Reactome, KEGG) and one database that contains expert curated disease associated genes (DisGeNET) (**Supplementary Table 3, Methods)**. Gene set analysis yielded 163 gene-set-trait associations with 14 unique gene sets (**Supplementary Table 4**). Given that 143 gene-set-trait associations were with 13 gene sets that included two genes with a well-established role in lipid biology (*APOB* and *APOC3*), we repeated the test removing variants in these genes. After removal, there was residual evidence of association (*p*<0.05) in 102 of 143 gene-set-trait signals representing 12 of 13 gene sets. Of the 163 gene-set-trait associations, the remaining 20 gene-set-trait associations (in gene sets not containing either *APOB* or *APOC3*) represent associations of various lipoprotein-related metabolic biomarkers with the “regulation of pyruvate dehydrogenase (PDH) complex” pathway in REACTOME (R-HSA-204174, min *p*= 7.85 × 10^−7^, trait=phospholipids in intermediate density lipoproteins (IDL-PL), **Supplementary Table 4**). These associations encompassed 12 LoF variants in WES and four in WGS (**Figure 1**). Upon further inspection, we found that most variants in this pathway were contributing to the association suggesting the signal was not driven by a single gene, in addition they all have the same direction of effect (i.e. the rho(ρ) value in the SKAT-O test was one in both the WES and the WGS analyses, **Supplementary Table 5**). Two variants were of particular interest as they were present in both WES and WGS datasets, rs113309941 in Pyruvate Dehydrogenase Complex Component X (*PDHX*) and rs201013643 in Pyruvate Dehydrogenase Phosphatase Regulatory Subunit (*PDPR*). In *PDHX*, rs113309941 leads to a premature stop mutation (Gln248Ter). It has an allele count (AC) of one in each of WES and WGS, and is very rare in the Genome Aggregation Database (gnomAD) (AC=3, allele number (AN)=246,116).rs201013643 in PDRP also leads to a premature stop (Arg714Ter) and is present in a single heterozygous individual in the WES dataset and two heterozygous in the WGS. This variant is also rare in gnomAD (AC=141, AN=275,988). The five participants carrying these two variants had higher than average values for biomarkers including cholesterol in intermediate-density lipoproteins (IDL-C) and low-density lipoproteins (LDL-C) (lying in upper percentile range from 44.1% to 0.03% for both traits), suggesting these variants may have a deleterious impact on lipid metabolism. Notably, one of the carriers of the *PDHX* Gln248Ter variant was in the top 0.03% for LDL-C in INTERVAL (4.086 mmol/l, 158.005 mg/dl) and had no predicted deleterious missense mutations in known hypercholesterolemia genes *PCSK9*, *APOB* or *LDLR* suggesting this novel protein-truncating variant may be a genetic cause for their high LDL-C levels. The other carrier was in the top 19.3% percentile of the cohort. None of the genes in this pathway has been previously associated with these traits and therefore this study links these genes collectively to IDL and LDL metabolism and circulating cholesterol for the first time.

**Figure 1-.**
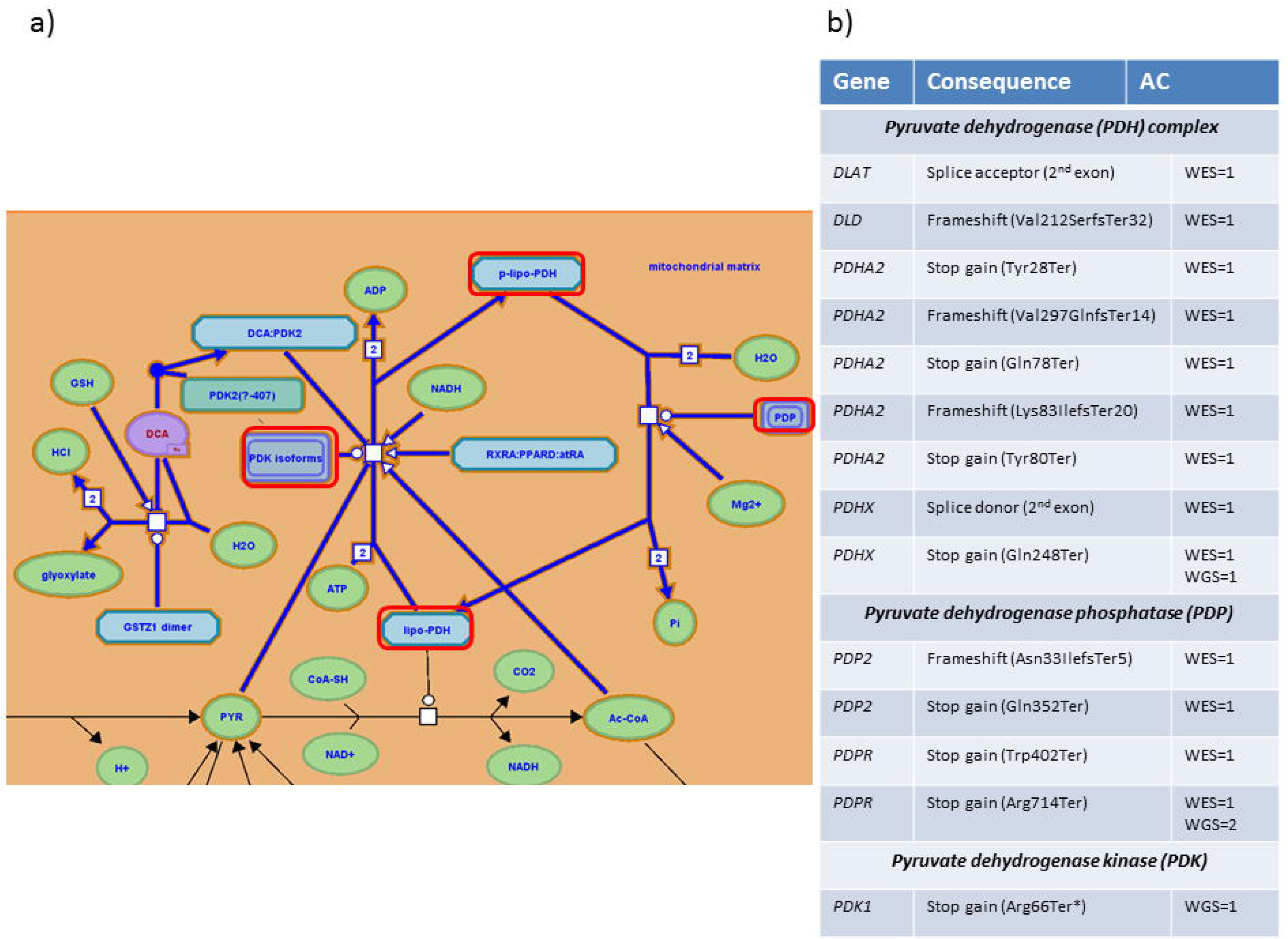
Loss-of-function (LoF) variants in regulation of pyruvate dehydrogenase (PDH) complex pathway. a) Figure adapted from REACTOME pathway browser [71]. Highlighted in red are protein complexes that carry LoF variants in INTERVAL WES or WGS. b) List of genes, consequences and allele count (AC) of LoF variants in the different protein complexes in the pathway.

### Enrichment of rare variant associations in genes near established GWAS signals in lipoprotein related metabolic biomarkers

Next, we conducted analyses to investigate whether genes near GWAS index variants associated with traditional lipid traits (HDL-C, LDL-C, TC and TG) were enriched for rare variant associations with high resolution lipoprotein measurements, which could suggest enrichment of effector transcripts in the gene set. Given that this was a hypothesis-driven approach using established signals, to boost discovery power we pooled together both WES and WGS data into a single dataset of 7,179 individuals. First, we extracted from the GWAS catalog (release 27-09-2017) the “reported genes” near signals that have been associated with HDL-C, LDL-C, TC or TG and created four gene sets (**Supplementary Table 6**). We only focused on genes that were reported unambiguously (i.e. where only one gene is reported) since for associations where more than one gene is reported, it is possible that only one will be the effector gene and rare variants from the non-effector genes will only add noise to the analysis and therefore reduce power. We grouped rare variants in the gene set using two nested approaches (LoF and MCAP+LoF) and ran SKAT-O on the gene sets for 157 lipoprotein and lipid traits. Using this approach we found associations (*p* < 0.005 correcting for effective number of tests, **Methods**) for genes near HDL GWAS signals with 18 HDL-related traits (**Supplementary Table 7**), the strongest association being with esterified cholesterol in extra-large HDL (XL-HDL-CE, *p*=2.83 × 10^−5^, MCAP+LoF). Associations (*p* < 0.005, **Methods**) in two extra-large HDL-C related traits remained after removing variants in genes known to be involved in conditions leading to abnormal lipid levels or genes where functional work has shown an effect on HDL-C (**Supplementary Table 8, Methods**) suggesting there is a contribution to the phenotypic variance of these traits by rare coding variants in genes, near GWAS signals, without a known role in HDL metabolism, which may represent novel effector transcripts.

### Enrichment of rare variation in tails of the phenotypic distribution of lipoprotein and glyceride related traits

Finally, we aimed to investigate whether individuals at the extreme tails of the phenotype distribution for 106 lipoprotein and lipid traits harboured rare coding variants likely to be contributing to their phenotype. We used the WES dataset as a discovery dataset and the WGS dataset as validation. An arbitrary cut-off of 10 individuals at each tail was used to define the tails for all of the 106 traits (**Methods**). After meta-analysis, we found an enrichment of deleterious rare variation (*p* < 0.00037, **Methods, Table 2, Supplementary Table 9**) in hyperlipidaemia related genes in the lower tail of cholesterol in small VLDL (S-VLDL-C), esterified cholesterol in small VLDL (S-VLDL-CE) and concentration of extra small VLDL particles (XS-VLDL-P), and rare variation in HDL remodelling related genes in the lower tail of concentration of small HDL particles (S-HDL-P). We still observed nominal evidence of association in the WES and WGS datasets for the S-VLDL-C and XS-VLDL-P results using a 0.5% percentile cut-off for the tails but no evidence of association was found when using a 1% percentile cut-off (**Supplementary Table 10**). This is likely due to the fact that by extending the number of individuals taken from the tails, we are decreasing the average distance to the mean and diluting signal coming from true extreme values.

**Table 2-.** Gene sets where there is a nominally significant enrichment of rare variation in the tails of a lipid or lipoprotein measurement (*p*>0.05) in both WES and WGS. Highlighted in bold are gene sets that are significant after meta-analysis using Stouffer’s method [69] and after adjusting for multiple traits (*p*<=0.00037). WES P: permutation *p* in WES. WGS P: permutation *p* in WGS. Meta-P: p after meta-analysis using Stouffer’s method. S-VLDL-FC: Free cholesterol in small VLDL. XS-VLDL-C: Cholesterol in very small VLDL. S-VLDL-C: Cholesterol in small VLDL. XS-VLDL-P: Concentration of very small VLDL particles. S-VLDL-CE: Cholesterol esters in small VLDL. S-HDL-P: Concentration of small HDL particles.

| Upper tails |  |  |  |  |
| --- | --- | --- | --- | --- |
| Trait | WES P | WGS P | Meta-P | Gene Set |
| S-VLDL-FC | $3.3 \times 10^{-2}$ | $2.37 \times 10^{-2}$ | $3.45 \times 10^{-3}$ | Hypertriglyceridemia_HPO |
| XS-VLDL-C | $3.3 \times 10^{-2}$ | $2.37 \times 10^{-2}$ | $3.45 \times 10^{-3}$ | Hypertriglyceridemia_HPO |
| Lower tails |  |  |  |  |
| trait | WES P | WGS P | Meta-P | Gene Set |
| S-VLDL-C | $5.8 \times 10^{-3}$ | $2.31 \times 10^{-3}$ | $7.61 \times 10^{-5}$ | Hyperlipidaemia |
| XS-VLDL-P | $1.85 \times 10^{-2}$ | $7 \times 10^{-4}$ | $9.42 \times 10^{-5}$ | Hyperlipidaemia |
| S-VLDL-CE | $5.8 \times 10^{-3}$ | $6.75 \times 10^{-3}$ | $2.07 \times 10^{-4}$ | Hyperlipidaemia |
| S-HDL-P | $2.72 \times 10^{-3}$ | $1.84 \times 10^{-2}$ | $2.89 \times 10^{-4}$ | HDL_remodeling |
| S-HDL-P | $4.10 \times 10^{-2}$ | $3.92 \times 10^{-2}$ | $8.24 \times 10^{-3}$ | Hypertriglyceridemia_CTD |

## Discussion

Exploring rare coding variation provides an opportunity to gain insights into biological processes regulating the circulating levels of metabolic biomarkers. Here we take advantage of the combination of sequencing data and high-resolution NMR measurements to elucidate how this variation influences multiple metabolic measurements in a healthy cohort of UK blood donors.

To identify genes and gene sets associated with metabolic biomarkers, we used a two-stage gene-based analysis using WES data in discovery (N_discovery_=3,741), and WGS data for validation (N_validation_=3,401). Rare-variant aggregation tests were used to identify genes harbouring multiple rare coding variants associated with metabolic biomarkers. To gain power at the validation stage we adjusted analyses for correlated traits, an approach previously described for single-point analysis [31]. This yielded significant power gains, notably for the known association of *PAH* with phenylalanine levels, where adjusting for 71 phenotypically correlated traits resulted in a greater than 30-fold magnitude change in the statistical evidence of association after meta-analysis. Overall, this approach yielded 4,114 gene-trait associations taken forward for validation (p_discovery_ <5 × 10^−3^). After meta-analysis, besides recapitulating previous associations in eight known genes (*APOB*, *APOC3*, *PAH*, *HAL*, *PCSK9*, *ALDH1L1*, SCARB1 and *LIPC*, **Table 1**), this method also identified four genes (*ACSL1*, *MYCN*, *B4GALNT3*, *FBXO36*) that met standard gene-level significance (*p*<2.5 × 10^−6^, **Table 1**) in at least one gene-trait association test. Of these, *ACSL1* and *MYCN* have been previously linked to lipid metabolism [34-36], and therefore will merit additional follow-up.

*ACSL1*, which codes for long-chain-fatty-acid—CoA ligase 1, is the predominant isoform of *ACSL* in the liver. The gene was associated with concentration of IDL particles in this study (*p* = 1.82 × 10^−7^), and its deficiency in the liver has been shown to reduce synthesis of triglycerides and beta oxidation, and alter the fatty acid composition of major phospholipids [37]. An intronic variant (rs60780116) in *ACSL1* has been associated with T2D [38] and elevated expression of *ACSL1* has been shown to be an independent risk factor for acute myocardial infarction [39].

*MYCN* encodes N-myc proto-oncogene protein and its amplification can lead to tumorigenesis [40,41]. Previous animal studies have shown that inhibition of *MYCN* can lead to accumulation of intracellular lipid droplets in tumour cells [36]. Here we find association between *MYCN* and concentration of lipids, phospholipids and triglycerides in medium VLDL, total particle concentration of medium VLDL and triglycerides in small VLDL (min *p* = 6.20 × 10^−7^, Table 1, **Supplementary Table 2**).

The other two genes do not have any obvious link to lipid metabolism. *B4GALNT3* encodes beta-1,4-N-acetyl-galactosaminyl transferase 3. This protein mediates the N,N’- diacetyllactosediamine formation on gastric mucosa [42]. Mouse knockouts have been associated with abnormal tail movements, abnormal retinal pigmentation and increased circulating alkaline phosphatase levels[43] and variants near the gene have been associated with height and hip circumference adjusted for BMI in human GWAS [44,45]. *FBXO36* is a member of the F-box protein family. F-box proteins are known to be involved in protein ubiquitination [46]. Replication of these signals in additional studies would represent a novel link between these genes and lipid metabolism.

In gene set analysis, the “regulation of pyruvate dehydrogenase (PDH) complex” pathway was newly associated with 20 traits, mostly related to IDL and LDL lipoproteins. None of the genes in this pathway have been previously linked to any of these phenotypes, and our data suggest the signal arises from a cumulative effect of loss-of-function variants in different genes in the pathway (Figure 1), which represents a novel link between this pathway and lipoprotein metabolism. Of note, a carrier of a rare stop gain mutation (Gln248Ter) in *PDHX* had very high levels of LDL-C (4.1 mmol/l, top 0.03% of full INTERVAL cohort) with no other rare mutation in genes known to harbour rare mutations causative of hypercholesterolaemia (*PCSK9*, *APOB*, *LDLR*). The other carrier of this variant had slightly increased LDL-C levels but within normal clinical range (1.823 mmol/l, top 19.3% of the full INTERVAL cohort). Since we lack information on medication, specifically, lipid lowering medication, the degree to which this variant influences the observed LDL-C levels is difficult to assess. The PDH complex has been shown to be crucial for metabolic flexibility, i.e. the capacity to adjust fuel oxidation based on nutrient availability, which itself has been shown to play a role in cardiovascular disease [47].

In analyses aiming at identifying effector transcripts at established GWAS loci associated with traditional lipid measurements (HDL-C, LDL-C,TC and TG), we established that reported genes mapping near HDL-C associated loci were enriched for rare coding variants associated with multiple HDL-related measurements. The results remained significant (p<0.005) after removing genes known to be directly involved in HDL metabolism. This suggests that rare variants in this gene set contribute to variation in these traits, and that this gene set is enriched for additional effector transcripts, although common variants in the same haplotype as these rare variants could also account for some of the observed signal.

Finally, we showed that one can detect enrichment of rare variation in genes involved in lipoprotein metabolism in phenotypic extremes of some of these NMR measurements. Specifically, we showed enrichment of rare variants in hyperlipidaemia related genes in individuals with very low levels of cholesterol and esterified cholesterol in small VLDL, and very low levels of the total concentration of small VLDL particles. Enrichment of rare variants in HDL remodelling genes in individuals with very low levels of small HDL particles was also observed. Given that high levels of small HDL particles have been previously associated with higher incidence of ischemic stroke [48] some of these variants could have protective effects. These results are in agreement with previous work on LDL-C [23] and HDL-C [49] that show that common polygenic signals seem to have a higher impact on the higher extremes of lipid traits whereas there is evidence for a higher prevalence of rare variation on the lower extremes [49]. This is also expected since the INTERVAL cohort consists predominantly of healthy blood donors and therefore might be depleted of individuals with rare “damaging” variants.

Altogether, our results show that focusing on rare variation and deep metabolic phenotyping provides new insights into circulating metabolic biomarker biology. This argues for the expansion of deeper molecular phenotyping as part of large cohort sequencing efforts to gain further understanding of the role of rare coding variation on circulating metabolic biomarkers which may potentially lead to novel drug target discovery and/or provide additional genetic validation for specific targets.

## Supporting information

## Acknowledgements

Participants in the INTERVAL randomised controlled trial were recruited with the active collaboration of NHS Blood and Transplant England (www.nhsbt.nhs.uk), which has supported field work and other elements of the trial. DNA extraction and genotyping was co-funded by the National Institute of Health Research (NIHR), the NIHR BioResource (http://bioresource.nihr.ac.uk/) and the NIHR Cambridge Biomedical Research Centre (www.cambridge-brc.org.uk) [*]. NMR measurements in the INTERVAL study were funded by the European Commission Framework Programme 7 (HEALTH-F2-2012-279233). The academic coordinating centre for INTERVAL was supported by core funding from: NIHR Blood and Transplant Research Unit in Donor Health and Genomics (NIHR BTRU-2014-10024), UK Medical Research Council (MR/L003120/1), British Heart Foundation (RG/13/13/30194), and NIHR Cambridge BRC. A complete list of the investigators and contributors to the INTERVAL trial is provided in reference [50]. The academic coordinating centre would like to thank blood donor centre staff and blood donors for participating in the INTERVAL trial.

*The views expressed are those of the author(s) and not necessarily those of the NHS, the NIHR or the Department of Health and Social Care.

**Di Angelantonio E, Thompson SG, Kaptoge SK, Moore C, Walker M, Armitage J, Ouwehand WH, Roberts DJ, Danesh J, INTERVAL Trial Group. Efficiency and safety of varying the frequency of whole blood donation (INTERVAL): a randomised trial of 45 000 donors. Lancet. 2017 Nov 25;390(10110):2360-2371.

IB and EZ acknowledge funding from Wellcome (WT206194). FRM received funding from Consejo Nacional de Ciencia y Tecnología México (489672). COW received funding from the British Heart Foundation Cambridge Centre of Excellence, (RE/13/6/30180) and Homerton College, University of Cambridge. We thank the Human Genetics Informatics (HGI) team at the Wellcome Sanger Institute for their contribution. We would also like to thank Patrick Short for discussions.

## Methods

### Participants

The INTERVAL cohort consists of 50,000 predominantly healthy blood donors in the UK [50]. All individuals have been genotyped using the Affymetrix UK Biobank Axiom Array and imputed using a combined UK10K-1000G Phase III imputation panel [51]. A subset of 4,502 individuals was selected for whole-exome sequencing (WES) [52] and another subset of 3,762 was selected for whole-genome sequencing (WGS). There was an overlap of 54 individuals in both datasets.

### Sequencing and genotype calling

WES and WGS were performed at the Wellcome Sanger Institute (WSI) sequencing facility. For WES, sheared DNA was prepared for Illumina paired-end sequencing and enriched for target regions using Agilent’s SureSelect Human All Exon V5 capture technology (Agilent Technologies; Santa Clara, California, USA). The exome capture library preparation was sequenced using the Illumina HiSeq 2000 platform as paired-end 75 bp reads. Reads were aligned to the GRCh37 human reference genome using BWA (v0.5.10) [53]. GATK HaplotypeCaller v3.4 [54] was used for variant calling and recalibration. For WGS, sheared DNA was prepared for Illumina paired-end sequencing. Sequencing was performed using the Illumina HiSeq X platform as paired-end 75 bp reads. Reads were aligned to the GRCh38 human reference genome using mostly BWA (v.0.7.12) although a subset of samples was aligned with v.0.7.13 or v.0.7.15. GATK HaplotypeCaller v3.5 was used for variant calling and recalibration. We extracted coordinates from the VCF files that mapped to regions targeted in the WES. We then used custom scripts to transform coordinates of variants to GRCh37 human reference.

### Sample QC

For WES data we filtered out samples based on the following criteria: i) withdrawn consent; ii) estimated contamination >3% according to the software VerifyBamID [55]; iii) sex inferred from genetic data different from sex supplied; iv) non-European samples after manual inspection of clustering in 1000G principal component analysis (PCA) and choosing cutoffs on the first 2 PCs; v) heterozygosity outliers (samples +/- 3 SD away from the mean number of heterozygous counts); vi) non-reference homozygosity outliers (samples +/- 3 SD away from the mean number of non-reference homozygous counts); vii) outlier Ti/TV rates (transition to transversion ratio +/- 3 SD away from the mean ratio); viii) excess singletons (number of singleton variants >3 SD from the cohort mean). After quality control 4,070 WES samples were kept. For WGS data we filtered out samples based on the following criteria: i) estimated contamination >2% according to software VerifyBamID; ii) non-reference discordance (NRD) with genotype data on the same samples >4%; iii) population outliers from PCA (PC1>0 and minimum PC2); iv) heterozygosity outliers (samples +/- 3 SD away from the mean number of heterozygous counts); v) number of third-degree relatives (proportion IBD (PI_HAT) >0.125) > 18, vi) overlap with WES. After quality control 3,670 WGS samples were kept.

### Variant QC

For variants with MAF>1% we used the following thresholds to exclude variants: i) VQSR: 99.90% tranche for WES and 99% tranche for WGS; ii) missingness>3%; iii) HWE p<1 × 10-5. For variants with MAF≤1% the following thresholds were used: i) VQSR: 99.90% tranche for WES, 99% tranche for WGS SNPs and 90% tranche for WGS indels; ii) GQ: <20 for SNPs and <60 for indels; iii) DP <2; iv) AB>15 & <80 for heterozygous variants. After genotype-level QC (GQ,DP,AB) only variants with <3% missingness were kept. 1,716,946 variants were kept in the final WES release and 1,724,250 in the final WGS release.

### Phenotype QC

A total of 230 metabolic biomarkers were produced by the serum NMR metabolomics platform (Nightingale Health Ltd.) [56] on 46,097 samples in the INTERVAL cohort. Glucose, lactose, pyruvate and acetate were excluded initially due to unreliable measurements. Conjugated linoleic acid and conjugated linoleic acid to total fatty acid ratio were set to missing for 3585 samples showing signs of peroxidation. Creatinine levels were set to missing for 1993 samples with isopropyl alcohol signals. Glutamine levels were set to missing for 347 samples that showed signs of glutamine to glutamate degradation. Samples with more than 30% missingness or identified as EDTA plasma were removed. After this step, for each pair of related samples (PI_HAT>0.125) we kept only one, preferentially keeping samples with the lowest missingness in WES or lowest NRD in WGS. We then separately performed linear regression for WES and WGS adjusting for age, gender, centre, processing duration, month of donation and 10 PCs. Residuals from both linear regressions were rank inverse normal transformed prior to use as the outcome variables in all subsequent analyses. After this final step we kept 3,741 samples in the WES dataset and 3,420 samples in the WGS dataset.

### Gene-based analyses

Coding variant consequences were annotated with VEP [57] using Ensembl gene set version 75 for the hg19/GRCh37 human genome assembly. Loss-of–function (LoF) variants were annotated with a VEP plugin: LOFTEE (https://github.com/konradjk/loftee). M-CAP scores were downloaded and we extracted all missense variants with AC>=1 in the WES or WGS datasets [29]. Two different nested tests were used to group rare variants into testable gene units: predicted to be high confidence LoF by LOFTEE in any transcript of the gene, and the same LoF variants plus rare (MAF <1%) missense variants mapping to any transcript of the gene predicted to be likely deleterious by M-CAP (M-CAP score >0.025) (MCAP+LoF).

We performed rare-variant aggregation tests as implemented in the SKAT-O R package [58,59]. For the LoF tests, we performed a burden test (rho=1) whereas for the MCAP+LoF tests we used the optimal unified approach (method=“optimal.adj”). Genes were taken forward for validation if *p*<5 × 10^−3^.

Adjusting for correlated phenotypes can increase power in single point association analyses [31], therefore to increase power, we implemented a strategy to incorporate information from the multiple phenotypes measured in our dataset. To minimise chances of a false positive association we only adjusted for phenotypes as covariates at the validation stage ensuring evidence of association in discovery stage was present without adjustment for covariates. In order for a metabolic biomarker to be selected as a covariate in the validation stage, the following conditions had to be met: i) no evidence of genetic correlation (*p*>0.05) with outcome using publicly available summary statistics from Kettunen *et al* (2016) [25]; ii) phenotypic correlation in our dataset >10%; iii) not belonging to same metabolic biomarker supergroup as outcome (**Supplementary Table 11**). This approach resulted in 99 eligible NMR traits for which other traits could be used as covariates. METASKAT [60] was used to perform meta-analysis using the same parameters as in discovery. A signal was considered to replicate if: i) it met our Bonferroni corrected gene-level significance threshold (*p* < 1.32 × 10^−7^); ii) >2 variants were tested; iii) it was nominally significant (*p* < 0.05) in the unadjusted test for WGS (i.e. without adjusting for correlated traits). The Bonferroni corrected gene-level significance threshold was chosen after adjusting the standard gene-level significance threshold (2.5 × 10^−6^) for 19 PCs explaining >95% of the variance of 226 metabolic biomarkers, an approach previously used in similar studies using the same NMR platform [24,25].

To test if a single variant was driving an observed association, we performed leave-one-out analysis for all variants contributing to the test. An association was considered to be driven by a single variant if, after removing it the test resulted in a non-significant association (*p* > *0.05*).

### Gene set analyses

To perform gene set analysis we obtained a curated gene-disease list from DisGeNET [61,62] and gene lists of metabolic pathways from KEGG [63-65] and Reactome [66,67] (**Supplementary Table 3**). The gene-disease list obtained from DisGeNET, combines expert curated gene-disease associations from the following databases: a) CTD (Comparative Toxicogenomics Database); b) UNIPROT; c) ORPHANET (an online rare disease and orphan drug data base); d) PSYGENET (Psychiatric disorders Gene association NETwork); and e) HPO (Human Phenotype Ontology). We limited analysis to gene sets with more than three genes. Finally we extracted loss-of-function variants from genes in the gene sets and ran SKAT-O (method=“optimal.adj”) for each of the traits. Similarly to the gene-based analysis, we used WES data as discovery, and took signals forward for validation in WGS if *p* < 0.01. Covariate selection for correlated traits was performed as described in the gene-based analysis. The Gene-set-wide significance threshold was calculated by first estimating the effective number of gene sets tested given the high overlap amongst them. Using PCA we estimated that 1094 PCs explain > 95% of the variance in gene sets. The significance threshold was therefore calculated as: 0.05/(1094*19)=2.41 × 10^−6^ where 19 corresponds to the effective number of phenotypes tested as described above. A signal was considered to replicate if after meta-analysis: i) it met our gene-set-wide significance threshold (*p*_*meta*_ < 2.41 × 10^−6^); ii) >2 variants were tested; iii) it was nominally significant (p_validation_<0.05) in the unadjusted test for WGS (i.e without adjusting for correlated traits).

### Genes near GWAS signals

GWAS catalog data files (release 27-09-2017) were downloaded from https://www.ebi.ac.uk/gwas/docs/file-downloads [68]. We focused on GWAS loci associated with HDL-C, LDL-C, TC and TG. We extracted all reported genes for GWAS loci that were associated at genome-wide significance (*p*<5 × 10^−8^) excluding cases where the “REPORTED GENE” value was: i) NR (not reported); ii) intergenic; iii) APO(APOE) cluster; iv) HLA-area (**Supplementary Table 6**). For this analysis, we ran SKAT-O using the optimal unified approach (method=“optimal.adj”) on the four gene sets (HDLC reported, LDLC reported, TC reported, TG reported, **Supplementary Table 6**). The list of genes known to be involved in conditions leading to abnormal lipid levels was created extracting relevant genes from the DisGeNET and Reactome gene lists. Afterwards, we conducted a manual review of the published literature to remove genes where functional work in mouse or human has revealed a direct role of the gene in HDL metabolism (**Supplementary Table 6**). The search terms used were “[gene name] loss of function HDL” and “[gene name] knockout HDL”. Significance threshold (*p* < 0.005) was determined by correcting for 10 PCs explaining >95% of the variance of the traits used in this analysis.

### Tails analysis

For this analysis, we used all lipoprotein and lipid traits but excluded derived measures (lipid ratios) resulting in 106 traits (**Supplementary Table 11**). We focused on likely deleterious missense and loss-of-function variation in lipid metabolism and disease gene sets (**Supplementary Table 12**) with an allele count <10 in each dataset. We chose an arbitrary cut-off of 10 individuals with the highest and lowest values for the traits to define our tails for all 106 traits. Given the high phenotypic correlation of our traits, there was a high overlap of individuals at the tails of the distributions (**Supplementary Figures 1-2**) so we removed traits that shared >=8 individuals with any other trait reducing the number of tested traits to 50. For each trait, total deleterious allele count from each gene set for upper and lower tails was obtained and an empirical *p* was calculated by performing 10,000 permutations extracting 10 random individuals from the phenotype distribution and counting the number of deleterious alleles from the gene set. The significance threshold (p = 0.00037) was chosen by correcting for 9 PCs explaining >95% of the traits variance and 15 pathways. Meta-analysis was done using Stouffer’s method [69] as implemented in the metap package [70] in R.

