## Supplementary material for "The influence of rare variants in circulating metabolic biomarkers"

^1^Wellcome Sanger Institute, Cambridge, UK, ^2^MRC/BHF Cardiovascular Epidemiology Unit, Department of Public Health and Primary Care, University of Cambridge, Cambridge, UK., ^3^The National Institute for Health Research Blood and Transplant Unit (NIHR BTRU) in Donor Health and Genomics, Department of Public Health and Primary Care, University of Cambridge, Cambridge, UK., ^4^Homerton College, Hills Road, Cambridge, CB2 8PH, UK, ^5^Department of Haematology, University of Cambridge, Cambridge Biomedical Campus, Long Road, Cambridge CB2 0PT, UK, ^7^NHS Blood and Transplant, Cambridge Biomedical Campus, Long Road, Cambridge CB2 0PT, UK, ^8^NHS Blood and Transplant - Oxford Centre, Level 2, John Radcliffe Hospital, Headley Way, Oxford OX3 9BQ, UK, ^9^Radcliffe Department of Medicine, University of Oxford, John Radcliffe Hospital, Headley Way, Oxford OX3 9DU, UK, ^10^University of Cambridge Metabolic Research Laboratories and NIHR Cambridge Biomedical Research Centre, Wellcome Trust-MRC Institute of Metabolic Science, Addenbrooke's Hospital, Cambridge, UK.

### Supplementary Figures

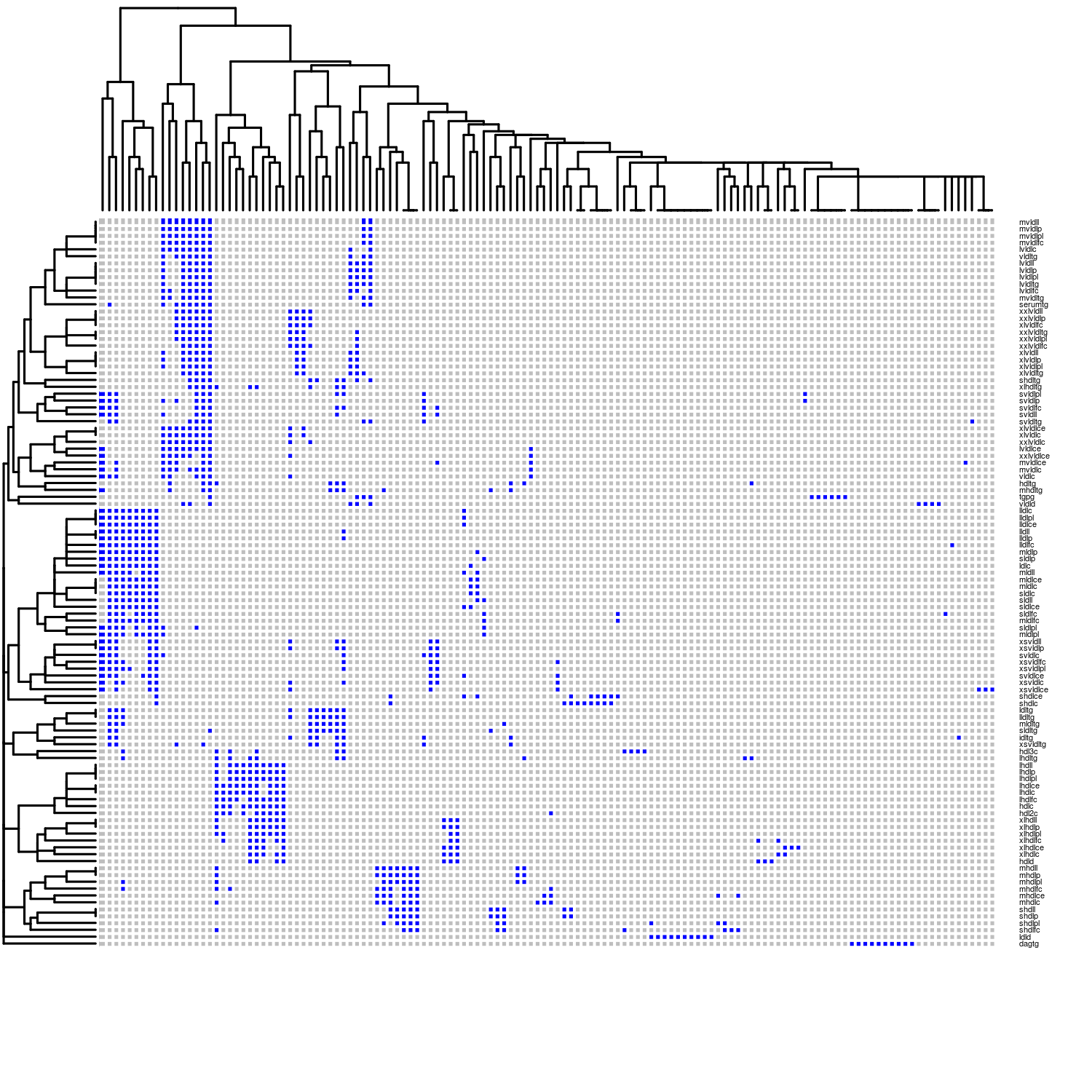

Supplementary Figure 1 – Overlap of top 10 participants in the tails of 106 lipid and lipoprotein traits. Columns represent the top 10 participants of at least one trait. Rows represent the 106 lipid and lipoprotein traits used in this analysis. A blue square represents presence of a participant in the top 10 participants for its respective trait.

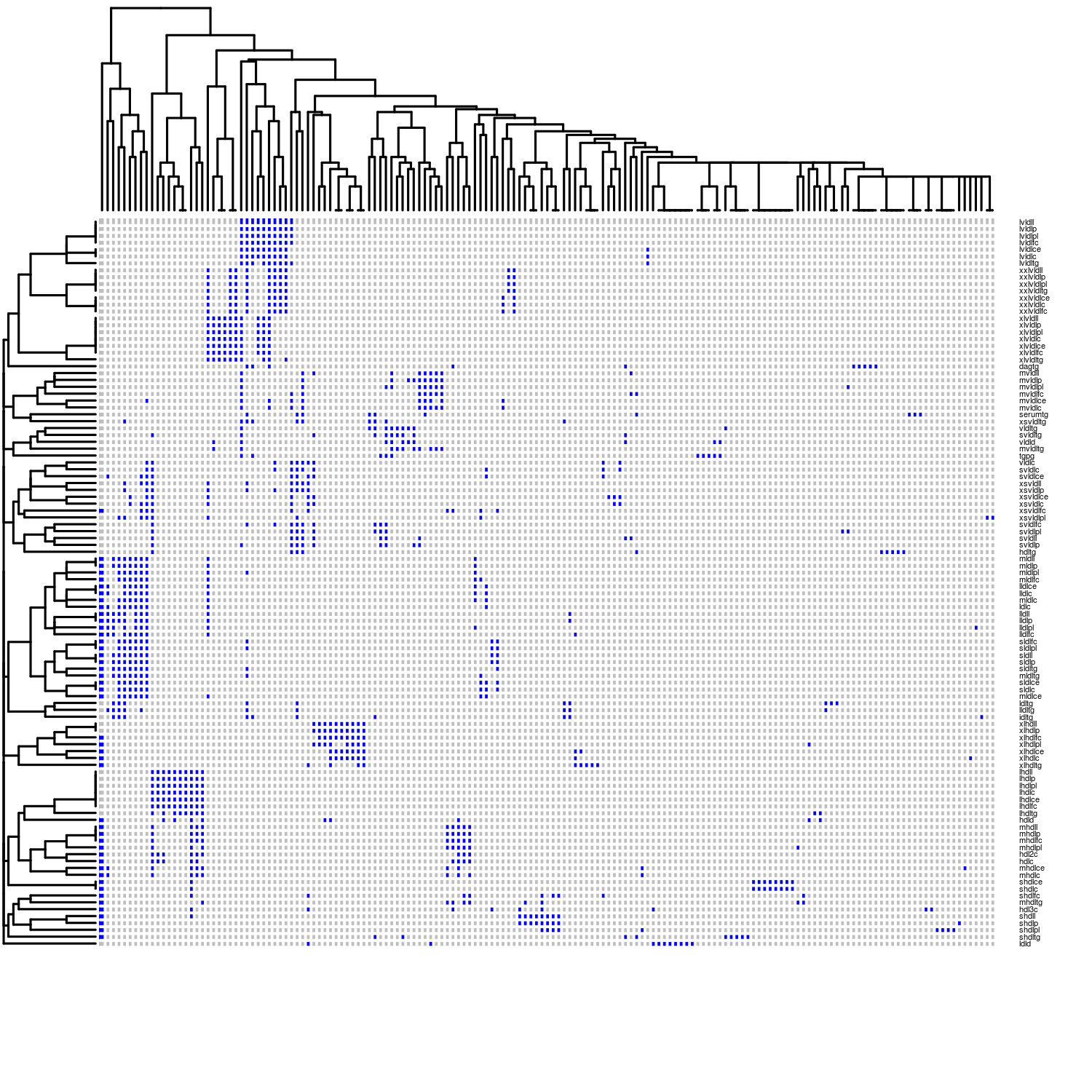

Supplementary Figure 2 – Overlap of bottom 10 participants in the tails of 106 lipid and lipoprotein traits. Columns represent the lower 10 participants of at least one trait. Rows represent the 106 lipid and lipoprotein traits used in this analysis. A blue square represents presence of a participant in the lower 10 participants for its respective trait.
